# *E. coli* translation strategies differ across nutrient conditions

**DOI:** 10.1101/224204

**Authors:** Sophia Hsin-Jung Li, Zhiyuan Li, Junyoung O. Park, Christopher G. King, Joshua D. Rabinowitz, Ned S. Wingreen, Zemer Gitai

## Abstract

For cells to grow faster they must increase their protein production rate. Microorganisms have traditionally been thought to accomplish this increase by producing more ribosomes to enhance protein synthesis capacity, leading to the linear relationship between ribosome level and growth rate observed under most growth conditions previously examined. Past studies have suggested that this linear relationship represents an optimal resource allocation strategy for each growth rate, independent of any specific nutrient state. Here we investigate protein production strategies in continuous cultures limited for carbon, nitrogen, and phosphate, which differentially impact substrate supply for protein versus nucleic acid metabolism. Unexpectedly, we find that at slow growth rates, *E. coli* achieves the same protein production rate using three different strategies under the three different nutrient limitations. Upon phosphate (P) limitation, translation is slow due to a particularly low abundance of ribosomes, which are RNA-rich and thus particularly costly for phosphorous-limited cells. In nitrogen (N) limitation, translation is slowed by limited glutamine and stalling at glutamine codons, resulting is slow elongation. In carbon (C) limitation, translation is slowed by accumulation of inactive ribosomes not bound to mRNA. These extra ribosomes enable rapid growth acceleration upon nutrient upshift. Thus, bacteria tune ribosome usage across different limiting nutrients to enable balanced nutrient-limited growth while also preparing for future nutrient upshifts.

## Introduction

Resource allocation during growth is a fundamental challenge faced by all cells^1-4^. For example, with a fixed resource budget, cells must balance production of the machinery that makes proteins (ribosomes, tRNAs, translation factors) with the production of the proteins themselves. This balance is generally represented by the RNA:protein ratio (R/P ratio)^5^. The R/P ratio captures protein production capacity, as >95% of total RNA is devoted to translation (rRNAs and tRNAs^5,6^). In single-cell organisms like *E. coli*, previous studies demonstrated that there is a linear relationship between R/P ratio and growth rate, with faster growth rates requiring more protein production capacity and therefore higher R/P ratios^5,7,8^. Production of ribosomes is costly as each contains 52 protein subunits and three large rRNAs^9,10^; hence, it is advantageous for the cell to saturate ribosomes with substrates. In this efficient ribosome scenario, the ribosome level should be fixed and independent of nutrient conditions for any growth rate, with the only way to increase protein synthesis rate being to increase the number of ribosomes^1,5,11^. One surprise for such a seemingly optimized system is that multiple studies have demonstrated that at slow growth rates *E. coli* accumulates inactive ribosomes^12^. There are two possible explanations for the presence of inactive ribosomes. First, it is possible that *E. coli* translation is constrained in such a way that it cannot function when ribosome levels drop too low^12^. Alternatively, *E. coli* could regulate ribosome production independently of growth rate. Here we settle this debate by showing that *E. coli* ribosome production and usage differ across nutrient conditions.

## Results

### Phosphate-limited cells achieve the same growth rate with fewer ribosomes than Carbon-or Nitrogen-limited cells

To determine the generality of the relationship between growth rate and ribosome content, we examined how the R/P ratio changes as a function of growth upon different nutrient limitations. We measured R/P ratios in *E. coli* under glucose- (C, carbon), ammonia- (N, nitrogen), and phosphate- (P, phosphorous) limitations over a range of different growth rates in chemostats (Figure 1A). Surprisingly, P-limited cells consistently exhibited lower R/P ratios than C-limited or N-limited cells, with a roughly 2-fold difference at the slowest growth rate tested (0.1 h^−1^, Figure 1B). Measured protein levels were similar in all cells regardless of growth rate or nutrient limitation (Figure S1A-B), indicating that the changes in the R/P ratios reflect changes in ribosome availability.

**Figure 1.**
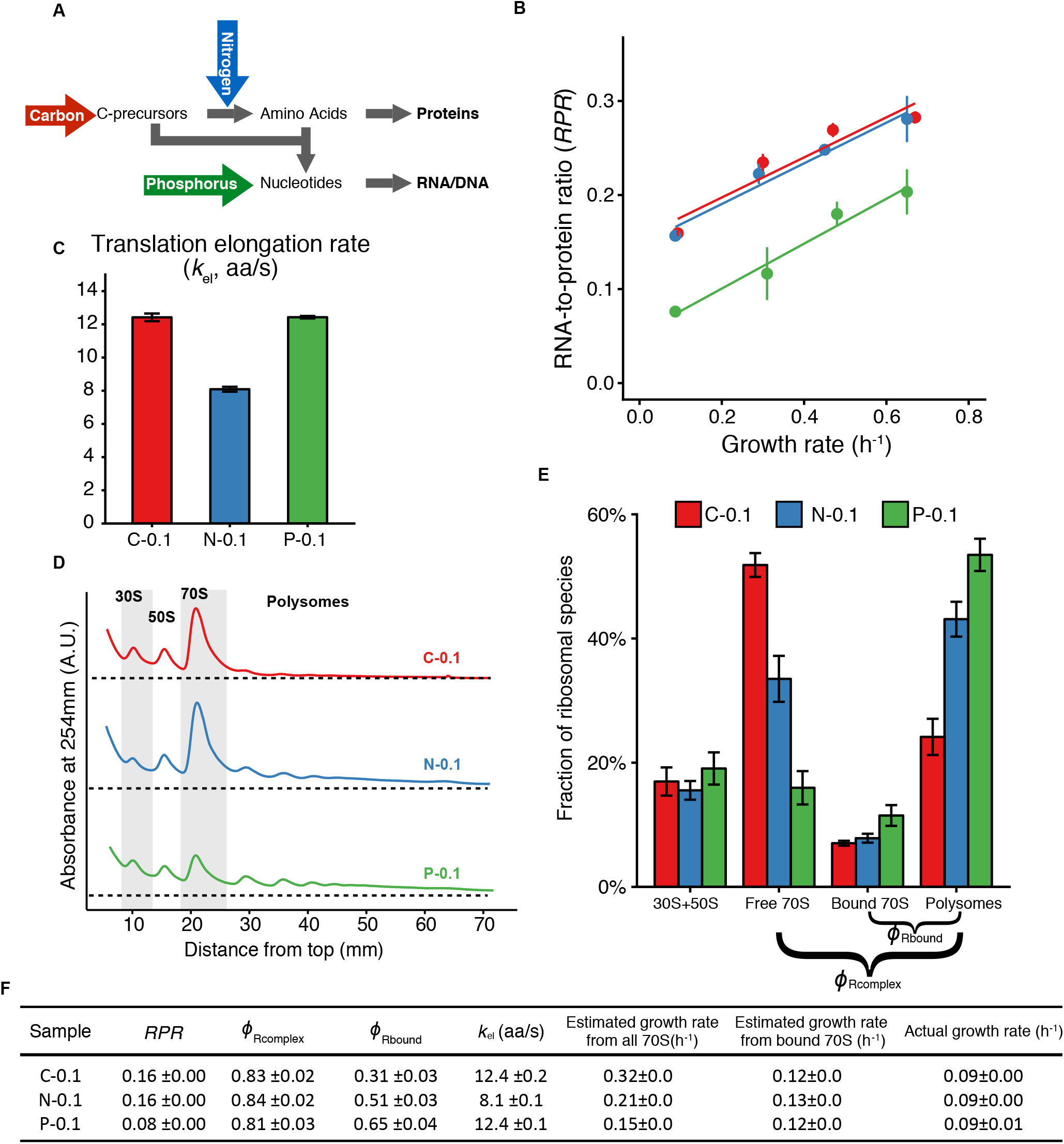
RNA-to-protein ratio is both growth-rate and nutrient dependent. **(A)** Schematic flow of nutrients for biomass formation. Carbon (C) and nitrogen (N) combine to make amino acids. Amino acids combine with carbon precursors and phosphorus (P) to make nucleic acids. **(B)** RNA-to-protein ratio for chemostat cultures upon C-, N-, and P-limitations at different growth rates. Each data point represents three technical replicates with standard deviations after error propagation shown as error bars. (See also Figure S1.) **(C)** Translation elongation rates (amino acids/s) as measured by the *lacZ* induction assay after correction for translation initiation. Error bars show the standard error from three biological replicates. **(D)** Polysome profiles of cells grown in C-, N-, P-limitations in chemostats at dilution rate 0.1 h^−1^. **(E)** Quantification of ribosomes in the form of subunits (30S + 50S), free 70S, mRNA-bound 70S (one ribosome on one mRNA), and polysomes (multiple ribosomes on one mRNA). Three biological replicates were analyzed with the error bars showing standard error of the mean. **(F)** Estimation of growth rate using measurements from RNA-to-protein ratio, elongation rates, and fractions of different ribosomal species.

The finding that *E. coli* can grow at the same rate with a lower R/P ratio implies that P-limited cells make the same amount of protein with fewer ribosomes, i.e. use ribosomes more efficiently. Thus, C/N-limited *E. coli* cells do not use ribosomes with optimal efficiency and their supply of “extra” ribosomes cannot be ascribed to requirements for productive translation. Since RNA accounts for two-thirds of the mass of bacterial ribosomes^9,14^, producing fewer ribosomes upon phosphate-limitation makes sense as a way for cells to deal with a limitation that preferentially reduces an elemental substrate needed to make RNA but not protein. This effect may be a direct consequence of low phosphate resulting in limited nucleotide pools and thus slow RNA synthesis, as deletion of phosphate sensing or storage systems, phoB^15^ or ppk^16^, did not alter the R/P ratio (Figure S1C-D).

### N-limited ribosomes translate slowly while C-limited cells accumulate more mRNA-free ribosomes

Why do C/N-limited cells accumulate so many ribosomes if P-limited cells can achieve the same protein synthesis rates with fewer ribosomes? One possibility is that the ribosomes in these cells translate slowly. We thus used a *lacZ* induction assay to compare the translation elongation rates of slow-growing C-, N-, and P-limited cells (0.1 h^−1^)^17,18^. We observed a reduced elongation rate in N-limited cells compared to C-and P-limited cells but no difference between C- and P-limited cells (Figure 1C). Thus, N-limited cells may need higher ribosome numbers to compensate for their slow translation elongation, but something else must explain the elevated ribosome numbers in C-limited cells.

To characterize ribosome pools we performed polysome profiling, which separates ribosome species using a sucrose gradient^19^. Regardless of the growth condition, all cells exhibited similar fractional pools of dissociated 30S and 50S subunits (Figure 1D). In contrast, the fraction of 70S monosomes was significantly larger in C/N-limited cells than in P-limited cells (Figure 1D). Since growth rate is proportional to protein synthesis rate, growth rate can be estimated by the product of the number of active ribosomes and the translation elongation rate. However, using the assumption that all 70S monosomes are active yielded very different growth rate estimates for C-, N-, and P-limited cells (Figure 1F and SI), which is inconsistent with the fact that these cells are growing at the same rate and have the same protein content. This inconsistency suggested that a fraction of the 70S ribosomes may not be active.

The mass of a single mRNA is small relative to the mass of a ribosome, such that 70S monosomes could represent either mRNAs with only one ribosome per transcript or inactive “free” ribosomes that are not associated with an mRNA. To distinguish free and mRNA-bound 70S monosomes, we utilized their differential sensitivity to high potassium levels (170 mM)^20^. High potassium causes free ribosomes to shift to a lower density but does not shift the density of mRNA-bound monosomes^20^. We thus designed a high-resolution “free-ribosome profiling” method to resolve this density shift (Figure S2). As controls, we confirmed that our assay detects the potassium-dependent shift of 70S monosomes induced by puromycin (Figure S2), which releases elongating ribosomes from their associated mRNAs^21^. There was no potassium-dependent shift detected in fast-growing cells that lack free ribosomes (Figure S2).

By combining traditional and free-ribosome polysome profiling, we quantified the relative fractions of all ribosome species in slow-growing C-, N-, and P-limited *E. coli*. The fraction of free monosomes was roughly 3-fold greater in the C-limited cells than in the P-limited cells, while the fraction of mRNA-bound monosomes remained relatively constant across nutrient limitations (Figure 1E). The accumulation of free 70S monosomes in C-and N-limited cells appears to be independent of a previously-described RaiA-dependent mechanism for ribosome storage as deletion of *raiA* had no impact on R/P ratios or polysome profiles (Figure S3). Importantly, revising our protein synthesis rate estimates to account for the fraction of inactive 70S monosomes yielded similar values for all cells, regardless of nutrient limitation (Figure 1F). These results both validate our experimental measurements and suggest that in different nutrient states, *E. coli* differentially tune ribosome number, translation rate, and active fraction to produce proteins at the same rates (Figure 2A).

**Figure 2.**
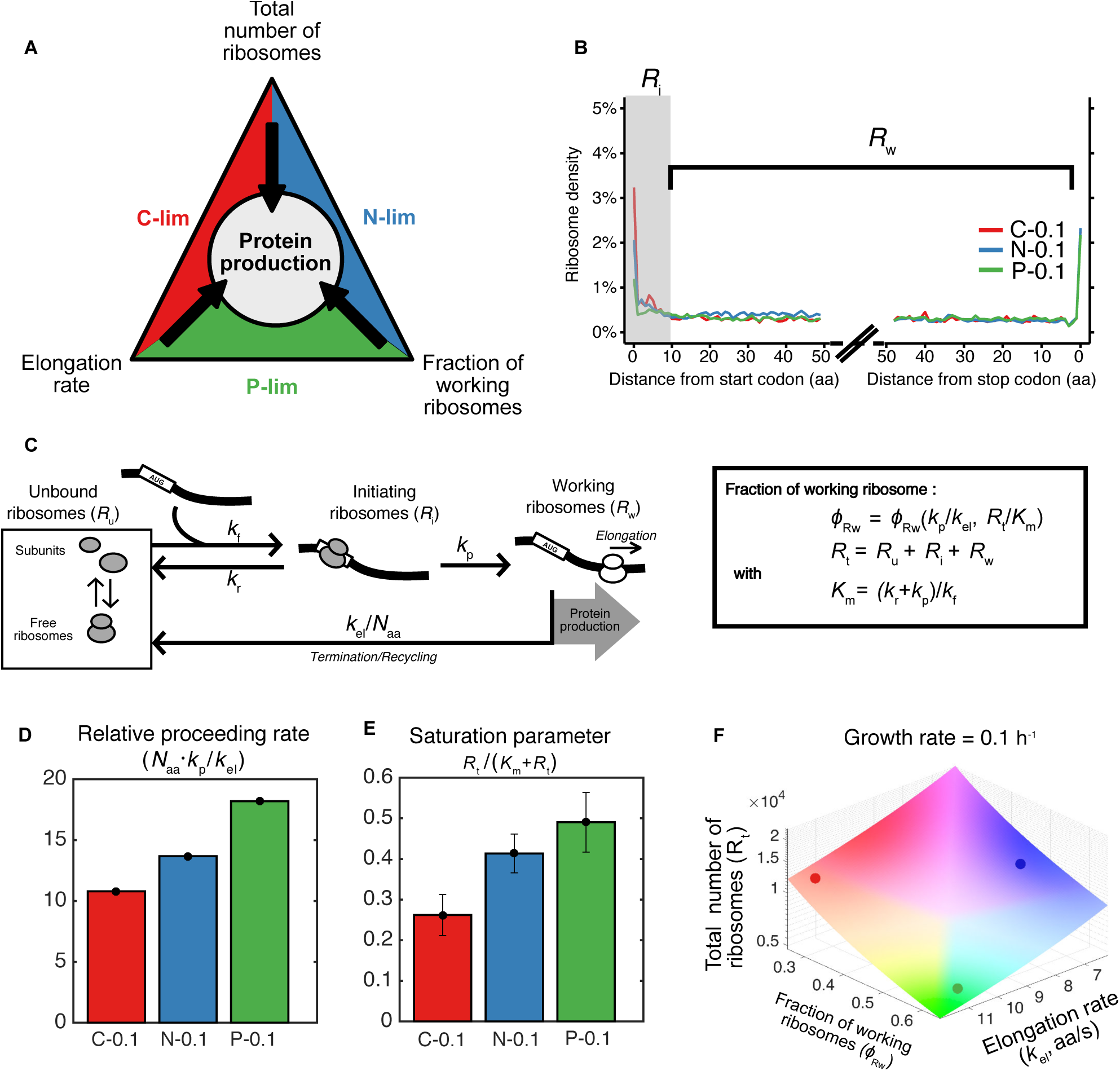
Macroscopic model reveals different ribosome dynamics that achieve the same growth rate. **(A)** Cells adapt to different nutrient limitations using different strategies of translational regulation that achieve the same protein production rate. **(B)** Averaged A-site ribosome counts within the first and last 50 codons of transcripts from ribosome profiling analysis. Ribosomes bound to the first 10 codons are defined as “initiating ribosomes” (*R*_i_) and those bound to the rest of transcripts except the stop codon are “working ribosomes” (*R*_w_). **(C)** Macroscopic model of ribosome dynamics. *k*_f_ forward rate (s^−1^); *k*_p_ proceeding rate (s^−1^); *k*_el_ elongation rate (aa/s); *N*_aa_ number of amino acids in an average protein of *E. coli*; *k*_r_ aborted translation rate (s^−1^); *R*_t_ total number of ribosomes. **(D)** The relative proceeding rate under C-,N-,P-limitations at growth rate 0.1 h^−1^. **(E)** The saturation parameter under C-,N-,P-limitations at growth rate 0.1 h^−1^. **(F)** The relationship between elongation rate (*k*_el_), fraction of working ribosomes (*ϕ*_Rw_) and total number of ribosomes (*R*_t_) that leads to the same growth rate at 0.1 h^−1^. Colored dots indicate the values for C-, N-, P-limited wild type cells.

### Quantitative modeling describes three different strategies of ribosome dynamics to achieve the same protein production rate

To better understand nutrient-dependent ribosome dynamics, we probed translation in slow-growing C-, N-, and P-limited cells by ribosome profiling. Analysis of ribosome densities as a function of distance from the start and stop codons revealed higher ribosome occupancy near the start codon (Figure 2B). Ribosome density thereafter was similar, with no decrease in ribosome density between the first and second halves of genes (Figures 2B and S4A), suggesting that there is little to no aborted translation after the first few codons. We quantified ribosome dynamics by building a simple macroscopic model (Figure 2C, S4B-E and details in SI). In this model, unbound ribosomes (*R*_u_) can bind to an mRNA with a free ribosomal binding site (with rate constant *k*_f_) to become initiating ribosomes (*R*_i_). The initiating ribosomes can proceed to elongation (with rate constant *k*_p_) to become working ribosomes (*R*_w_), or can abort translation (with rate constant *k*_r_). Working ribosomes elongate to finish translation and become unbound (with rate constant *k*_el_/*N*_aa_), where *N*_aa_ is the length of an average protein in amino acids. We defined the fraction of ribosomes bound to the first 10 codons as initiating ribosomes since the ribosome footprint size is ~10 codons^22,23^. To validate our model we used it to calculate elongation rates, which closely agreed with those we measured experimentally (Figure S5A).

Analysis of our model indicated that the system can be characterized by two main dimensionless parameters: the “relative proceeding rate”, defined as the ratio between the rate of ribosomes proceeding from initiation to elongation (*k*_p_, s^−1^) and the rate of elongation (*k*_el_ /*N*_aa_, s^−1^); and the “saturation parameter” (*R*_t_ /(*K*_m_+*R*_t_)), reflecting the degree of saturation of ribosome binding sites on mRNAs, where *R*_t_ is the total ribosome number and *K*_m_ = (*k*_r_ + *k*_p_)/*k*_f_ (Figures 2D-E and S5B-C). Fitting the measured ribosome densities, translation elongation rates, and pool sizes of ribosome species to the macroscopic model revealed that P-limited cells have the highest relative proceeding rate while C-limited cells have the lowest saturation parameter, and N-limited cells have intermediate values for both parameters (Figures 2D-E and S5B-H). Thus, C-, N-, and P-limited cells produce proteins at the same rate using three different strategies: P-limited cells have few ribosomes that are mostly active and elongate rapidly, N-limited cells have more ribosomes but fewer are active and they elongate slowly, and C-limited cells have many ribosomes, which elongate rapidly, but even fewer are bound to mRNA (Figure 2F and S5I).

### N-limited ribosomal regulation is mediated by RelA, the ppGpp alarmone synthase

Insight into the molecular basis of nutrient-specific ribosome regulation came from analysis of the codon occupancies in our ribosome profiling data. Codon-specific ribosome stalling leads to increased codon occupancy and is a hallmark of insufficient pools of the corresponding charged tRNAs. P-limited cells exhibited no elevated codon frequencies, consistent with these cells’ efficient ribosome usage (Figure 3A). In contrast, both C-and N-limited cells exhibited significant codon-specific stalling. Under N-limitation, ribosomes stalled at both of the two glutamine-encoding codons, which together account for 4.4% of all predicted ORF codons in *E. coli*. This result is consistent with previous studies indicating that glutamine is the most strongly-depleted amino acid pool upon N-limitation and serves as an intracellular sensor for extracellular nitrogen levels^24-26^. Upon C-limitation, we observed elevated occupancy of the Leu-CUA codon, which was surprising as there is no known intracellular carbon sensor that controls translation. *E. coli* has six leucine codons decoded by five leucine tRNA species^27,28^. Leu-CUA is the rarest Leu codon, accounting for only 0.4% of all predicted ORF codons.

**Figure 3.**
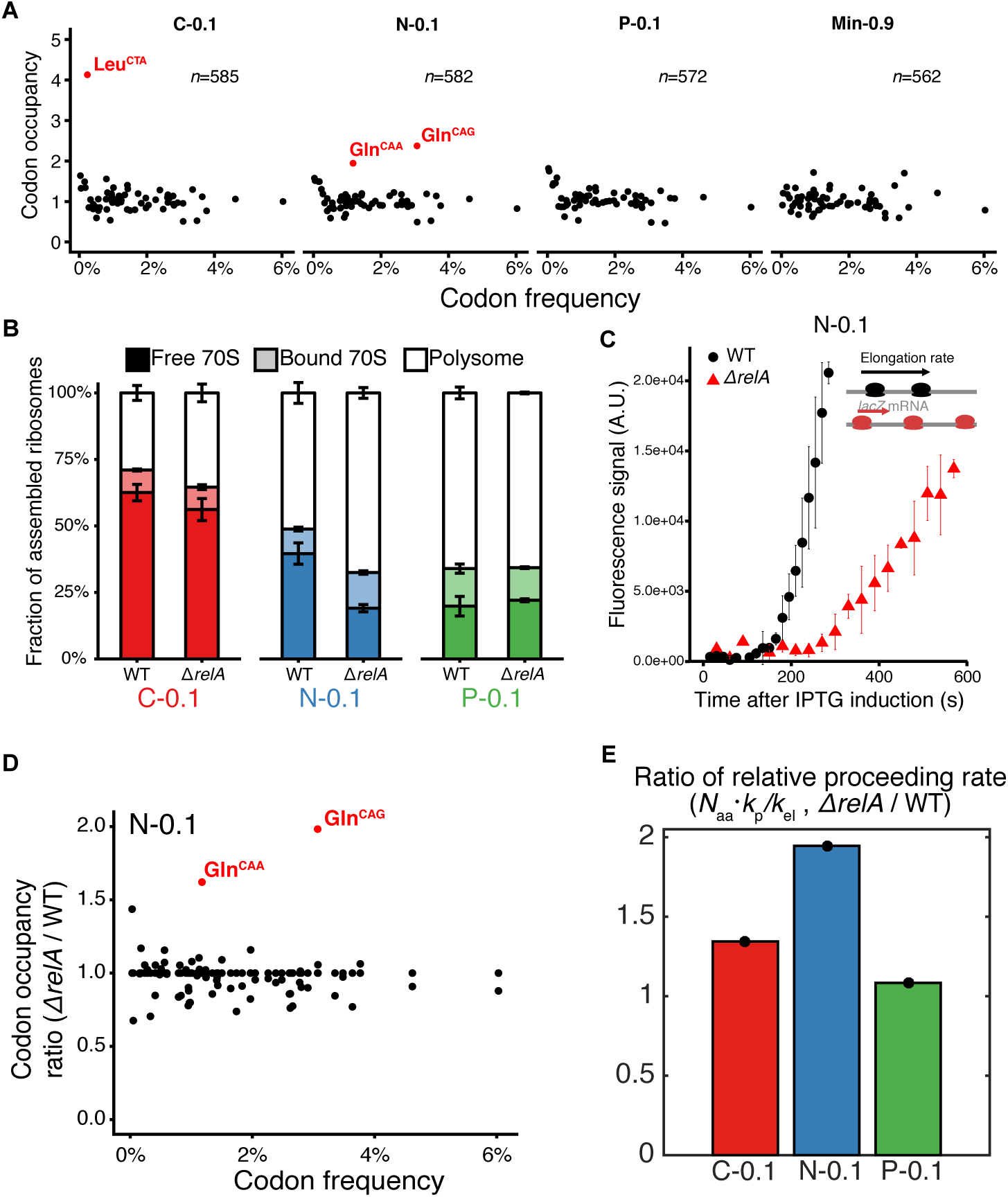
Deletion of *relA* disrupts translation regulation under nitrogen limitation. **(A)** A-site codon occupancy under different growth conditions from ribosome profiling. Occupancy was calculated as the ratio between measured and expected counts for each gene based on codon frequency. The average of this ratio is plotted (with number of genes specified). Codons that have higher than average ratio by 2.5 standard deviations are highlighted. **(B)** Fraction of assembled (70S) ribosomes in wild type and *∆relA* under C-, N-, and P-limitation at growth rate 0.1 h^−1^. Error bars show the standard error from three biological replicates. **(C)** *lacZ* induction assay for wild type and *∆relA* under nitrogen limitation at growth rate 0.1 h^−1^. The lag time measures when the first functional LacZ is produced and is inversely proportional to the elongation rate. The error bars show standard error from three biological replicates at each time point. **(D)** Ratio of codon occupancy between *∆relA* and wild type under nitrogen limitation at growth rate 0.1 h^−1^. (E) Ratio of the relative proceeding rate (*N*_aa_*·k*_p_/*k*_el_) between *∆relA* and wild type across different conditions at growth rate 0.1 h^−1^.

In addition to clarifying the link between metabolism and translation for N-limitation, the observation that ribosomes stall at specific codons upon C/N-limitation suggested a molecular mechanism for nutrient-specific translation regulation (Figure 3A). In bacteria, insufficient charged tRNA pools activate the stringent response to induce accumulation of the cellular alarmone, ppGpp^30,31^. ppGpp is known to regulate rRNA transcription *in vivo*. Its role in translation is less well-understood but ppGpp has been shown *in vitro* to inhibit translation factors such as EF-Tu and IF2 by competing with GTP^32,33^. To test how ppGpp accumulation might affect translation *in vivo*, we induced ppGpp synthesis by treating batch-grown *E. coli* with serine hydroxamate (SHX). SHX is a serine analog that competitively inhibits serine tRNA synthetase to yield uncharged serine-tRNA and thereby activate the RelA ppGpp synthase^34^. SHX treatment increased the pool of mRNA-free 70S monosomes, and this effect was completely dependent on *relA* (Figures S6A-B). Thus, inducing ppGpp by activating RelA alters translation by increasing the fraction of inactive free 70S ribosomes.

RelA is primarily required for the accumulation of ppGpp upon N-limitation but not C-limitation^31^. Consistently, deletion of *relA* had little impact on the accumulation of free ribosomes upon C-limitation, but significantly reduced free ribosome pools upon N-limitation to levels similar to those observed upon P-limitation (Figures 3B and S6C). We could not probe the role of ppGpp in C-limited ribosome accumulation because unlike N-limitation, C-limitation elevates ppGpp through *spoT*, which is essential^35^. N-limited *∆relA* cells also contained more polysomes than wild type (Figure 3B), suggesting that these cells had a higher fraction of elongating ribosomes. This result was initially confusing because the wild type and *∆relA* N-limited cells were grown at the same growth rate and maintained the same R/P ratio (Figure S6D). We thus measured the rate of translation elongation and found that while N-limited *∆relA* cells have higher fractions of translating ribosomes, their ribosomes elongate more slowly (Figure 3C), resulting in the same rate of protein production as wild type.

To understand how RelA influences ribosome dynamics we performed ribosome profiling. N-limited *∆relA* cells exhibited more pronounced ribosome stalling at both glutamine codons than wild type N-limited cells, while *relA* deletion had no effect on P-or C-limited cells (Figures 3D and S7). Fitting the *∆relA* cells measurements to our ribosome dynamics model revealed that *∆relA* cells specifically increase the relative proceeding rate under N-limitation, but display no effect on relative proceeding rate under P-or C-limitation and no effect on the saturation parameter in any condition (Figures 3E and S5B-C). These results suggest that, consistent with its known effects on IF-2 *in vitro*, in N-limited cells RelA functions to restrict the transition from translational initiation to elongation. Thus, in the absence of RelA, more ribosomes attempt to elongate, which exacerbates the depletion of charged tRNA pools, leading to increased stalling and a slower translational elongation rate.

### Extra ribosomes may facilitate growth acceleration upon nutrient upshift

While cells can modulate different aspects of ribosome dynamics to achieve the same protein production rate, what benefits might be served by the inefficient translation system used by C-limited cells where many ribosomes are inactive? Since free ribosomes accumulate the most at the slowest growth rates, we hypothesized that our findings could reflect a trade-off between steady-state growth rate and the ability to respond to a fluctuating environment. Such transitions could include rapidly and safely slowing growth when nutrients become depleted and rapidly increasing growth rate when nutrients are replenished. In this scenario, cells may benefit more by optimizing their ability to rapidly utilize new nutrients, for example to outcompete their neighbors or maximally utilize a transient pulse of nutrients, than by optimizing steady-state growth rate when nutrient levels are low.

We built a mathematical model of cellular resource allocation to predict the growth dynamics upon upshift. This model supported the hypothesis that the larger free ribosome pools of C/N-limited cells should enable them to increase their growth rates more quickly than P-limited cells upon nutrient upshift (Figures 4A, S8 and details in SI)^4,36^. We experimentally tested this prediction by measuring the growth rates of slow-growing (0.1 h^−1^) *E. coli* immediately after being shifted to rich media (LB + 0.4% glucose) ^37^. As predicted, C/N-limited cells increased their growth rates significantly faster than P-limited cells (Figure 4B). Thus, the distinct translation strategies employed under different nutrient conditions may represent nutrient-specific adaptations, with P-limited cells optimizing for current steady-state growth under slow growth conditions, and C-and N-limited cells favoring the ability to rapidly recover growth (Figure 4C).

**Figure 4.**
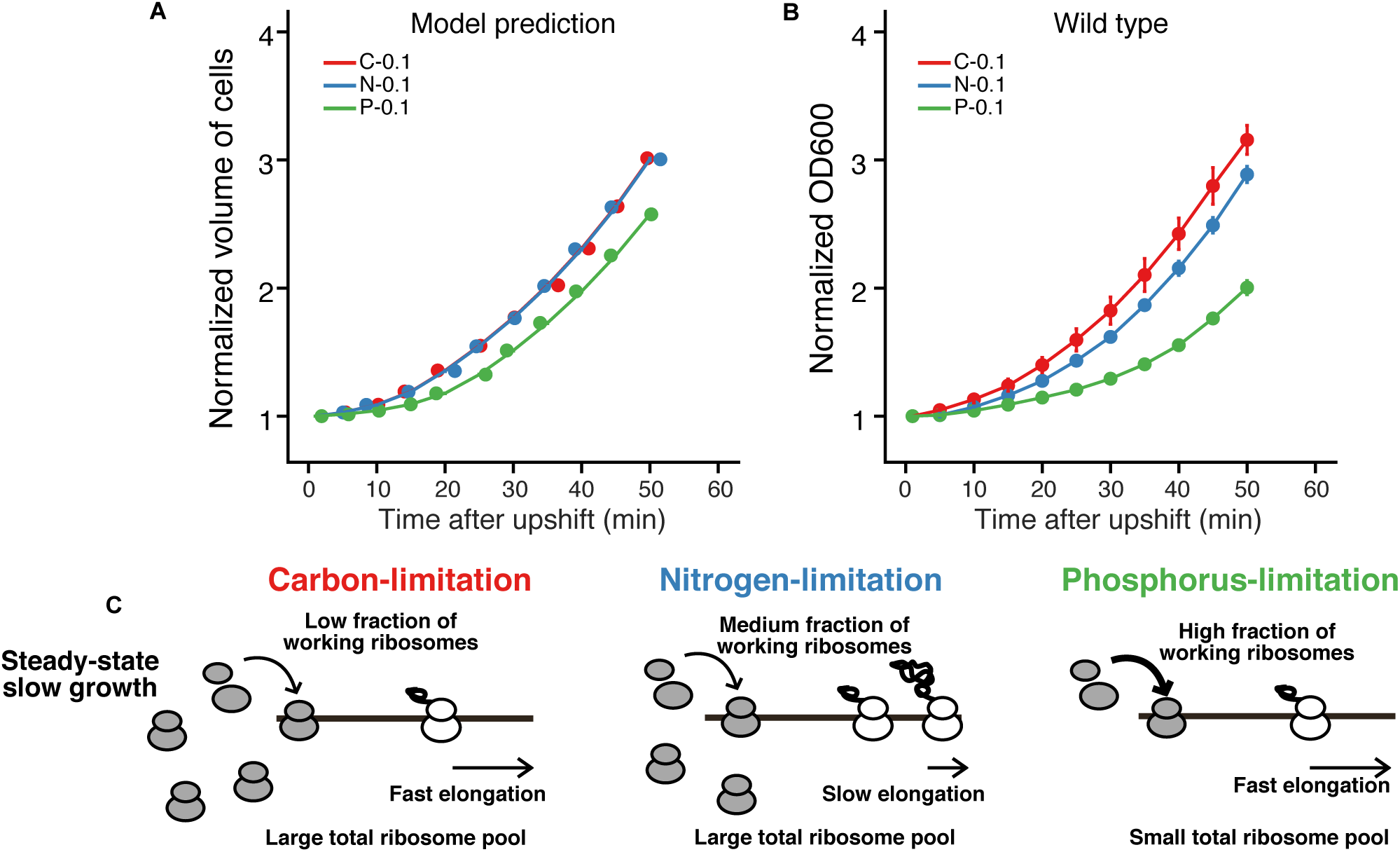
Extra ribosomes confer growth advantage upon nutrient upshift. **(A)** Theoretically predicted growth curves from the macroscopic model. **(B)** Experimental nutrient-upshift growth curves in LB + 0.4% glucose for wild type cells from C-, N-, and P-limited chemostats at growth rate 0.1 h^−1^. **(C)** Model for nutrient-dependent ribosome usage: Under C-or N-limitation, the total ribosome pool is high while under P-limitation the total pool is low. C-limited cells elongate fast but have a low fraction of working ribosomes. N-limited cells elongate slower but have a higher fraction of working ribosomes than C-limited cells. Under P-limitation the low supply of ribosomes leads to both a high fraction of working ribosomes and fast elongation to meet the protein production demand.

## Discussion

In previous studies, *E. coli* were found to vary R/P ratio with growth rate independently of the specific nutrient limitation used to produce a given growth rate^5,8^. Meanwhile, transcriptional analysis in *Saccharomyces cerevisiae* suggested that the primary determinant of the response to a wide range of stresses was the cellular growth rate rather than the specific stressor^38^. Together, these studies suggested that the primary regulator of microbial physiology is growth rate. However, our findings demonstrate that at the same growth rate, *E. coli* exhibit significantly different translation strategies across nutrient limitations; at the lowest growth rate tested, P-limited cells produced the same amount of protein with roughly half as many ribosomes as C/N-limited cells. P-limited cells also exhibited smaller inactive 70S monosome pools and higher relative proceeding rates than C/N-limited cells. Furthermore, while C/N-limited cells have similar ribosome numbers, they also display differences in free ribosome pools, translational elongation rates, and sensitivity to the loss of the ppGpp synthase RelA. Thus, our results suggest that the extra ribosomes of C/N-limited cells do not reflect essential constraints, but rather reflect a selectively beneficial adaptation.

Our findings also implicate ppGpp as a key mediator of ribosome activity that specifically modulates the transition from translation initiation to elongation. ppGpp is a well-characterized cellular alarmone that senses translational activity and inhibits rRNA transcription^31,39^. However, ppGpp can competitively inhibit GTP-dependent enzymes, including the translation initiation factor IF-2 that is required for the transition to elongation^40^. Our data thus provide *in vivo* support for a previous *in vitro* study demonstrating that ppGpp inhibits IF-2 function^33^. Since ppGpp can also affect other GTP-binding proteins involved in translation^32,41^, future studies on bacterial translation will confirm exactly how ppGpp interferes with ribosome function.

In native environments such as the mammalian gut, bacteria such as *E. coli* are faced with feast and famine cycles induced by feeding cycles^42^. Here we show that the linear relationship between R/P ratio and growth rate in *E. coli* does not reflect the optimization of steady-state growth rate but, instead, reflects the ability of cells with higher inactive ribosome pools to rapidly accelerate growth upon nutrient repletion. These findings suggest that *E. coli* may improve their fitness by sacrificing their maximal growth rate in nutrient-poor periods in return for the ability to respond to a changing environment, including rapidly accelerating growth in nutrient-rich periods. Our macroscopic model highlights how cells can tune total ribosome number, the fraction of working ribosomes, and the rate of translational elongation to achieve the same total protein production rate while balancing other constraints such as reduced amino acid availability or the need to rapidly accelerate growth. Future studies will address the consequences of adaptation strategies in dynamic conditions, for example the generality of bacteria optimizing growth rate transitions at the expense of steady-state growth^43^, but this strategy could explain recent reports of the sub-optimality of protein allocation for *E. coli* in the presence of poor carbon sources^44^, as well as the sub-optimal expression levels of essential genes in *B. subtilis*^45^.

## Author Contributions

S.H.L., J.O.P., J.D.R., N.S.W, and Z.G. designed the experiments. S.H.L., J.O.P., and C.K. performed experiments. Z.L. and N.S.W. constructed the mathematical models. S.H.L. conducted computational analysis of sequencing data. S.H.L, Z.L., N.S.W, and Z.G. wrote the paper with assistance from J.D.R.

## Acknowledgments

We thank the members of the Gitai, Wingreen, and Rabinowitz labs for helpful discussions. We thank Gene-Wei Li for his support for the ribosome profiling experiments. We thank the Microarray Core Facility at the Lewis-Sigler Institute (Daniel Sanchez, Jennifer M. Miller, Jessica Wiggins, and Wei Wang) for RNA-Seq sample processing and sequencing and the Princeton Proteomics Core Facility (Henry Shwe and Tharan Srikumar) for ribosome profiling sample processing. Lance Parsons provided helpful technical support for bioinformatics analysis of the sequencing data. We thank the Botstein lab, particularly Sandy Silverman, for chemostat operation support and all former members for discussions relating to microbial growth.

The ribosome sequencing data that support the findings of this paper will be deposited in GenBank with the accession code at publication.

This work was supported by grants from the NIH (DP1AI124669 and R01GM082938).

## Materials and Methods

### Cell strains and growth conditions

*Escherichia coli* strain NCM3722 was grown in batch or continuous cultures. To achieve different growth rates, different carbon or nitrogen sources were provided in batch culture, whereas dilution rates ranging from 0.1 h^−1^ to 0.7 h^−1^ were used in continuous (chemostat) cultures. The chemostat (Sixfors, HT) volume was 300mL with oxygen and pH probes to monitor the culture. pH was maintained at 7.2 +/− 0.1 and the aeration rate was set at 4.5 l/h. 40 mM MOPS media (M2120, Teknova) was used with glucose (0.4%, Sigma G8270), ammonia (9.5 mM NH_4_Cl, Sigma A9434) and phosphate (1.32 mM K_2_HPO_4_, Sigma P3786) added separately. For carbon-and nitrogen-limiting media, glucose and ammonia concentrations were reduced by 5-fold (0.08% and 1.9mM respectively). Phosphorus-limiting medium contains 0.132 mM K_2_HPO_4_. *∆relA* mutant was generated by P1 transduction from the KEIO collection^46^ into *Escherichia coli* strain NCM3722

### Nutrient upshift growth measurement

Cells from chemosats were mixed with 4X volumes of fresh pre-warmed media and grown in flasks in a 37°C water bath. Cell growth was monitored every 5 minutes by checking absorbance at 600nm in a quartz cuvette (Starna, 16.160-Q-10/Z8.5) using a spectrophotometer (GENESYS™ 20, Thermo Scientific). The LB media (244610, BD) used for upshift was supplemented with 0.4% glucose.

### Total RNA measurement

The method for RNA measurement was adapted from You *et al.*^1^. 1.5 mL of cultures were pelleted by centrifugation for 1 min at 13,000 X g. The pellet was frozen on dry ice and the supernatant was taken to measure absorbance at 600 nm for cell loss. The pellet was then washed twice with 0.6M HClO_4_ and digested with 0.3M KOH for 1 hour at 37°C. The solution was then precipitated with 3M HClO_4_ and the supernatant was collected. The pellet was re-extracted again with 0.5M HClO_4_. The supernatants were combined and absorbance measured at 260nm using NanoDrop (ND-1000, NanoDrop). Total RNA concentration was determined by multiplying the A260 absorbance with 31 (μg RNA/mL) as the extinction coefficient.

### Total protein measurement

The method for protein measurement is adapted from You *et al.*^1^. 1.5 mL of cell cultures were pelleted by centrifugation for 1 min at 13,000 X g. Cells were washed with 1 mL MOPS buffer once, re-suspended in 200 uL water and left on dry ice. All the supernatant was collected and OD600 was measured for cell loss. To measure the protein content, samples were thawed, 100μL 3M NaOH were added, and the sample was heated for 5 min at 98°C. The samples were cooled down to RT for 5 min before 300μL 0.1% CuSO_4_ were added for biuret assay. The samples were incubated at RT for 5 min and centrifuged at 13,000 X g for 1 min. The supernatant was collected and absorbance was measured at 555nm for a 200μL sample volume in a microplate reader (Synergy HT, BioTek) with software Gen 5.0. Proper dilution of albumin (23209, Thermo) with known concentrations was used to infer the total protein concentration in the cell.

### Polysome profiling and quantification of ribosome fraction

200mL of cells were collected from cultures by filtration through 90mm cellulose acetate membranes with a 0.2 μm pore size (CA029025, Strelitech) at 37°C, scratched with a clean and pre-warmed stainless steel spatula, and snap-frozen in liquid nitrogen. The whole filtration procedure did not take more than 2 minutes in order to maintain the original physiological state. Cell pellets were mixed with 650μL lysis buffer frozen nuggets (20mM Tris-HCl pH 8.0, 10mM MgCl_2_, 100mM NH_4_Cl, 0.4% Triton X-100, 0.1% NP-40, 1 mM Chloramphenicol, 100 U/mL RNase-free DNase I (04716728001 Roche)) in a pre-chilled 10mL jar (014620331, Retsch). Pulverization was done by cryomill (Retsch) at 15 Hz for 15 minutes. The thawed cell lysates were quantified by NanoDrop and 200μL of lysates with RNA concentration ranging from 80μg to 500μg were used. For overall polysome quantification, lysates were loaded to 10%-55% linear sucrose gradients (20mM Tris-HCl pH 8.0, 10mM MgCl_2_, 100mM NH_4_Cl, 300 μM Chloramphenicol) made by GradientMaster (BioComp). The gradients were placed in a SW41Ti bucket and centrifuged in Optima XE-100 Ultracentrifuge (Beckman Coulter) at 35,000 rpm for 2 hours at 4°C. Gradients were fractionated by BioComp Gradient Fractionator and the absorption curves at 254nm were recorded by a UV monitor (EM-1, BioRad).

Quantification of the polysome profiles was done using customized MATLAB codes. First, baselines were estimated using the readings where no peaks existed and the background was subtracted. The background from the free nucleotides and tRNA was removed by fitting an exponential decay function to the first peak representing the source of non-ribosome signals. Each ribosome peak was picked and quantified by integrating the area underneath the curve. To quantify different species of ribosomes in the 70S peak, 100mM NH_4_Cl was replaced with 170mM KCl. Cell lysates were loaded onto 10%-30% linear gradient and centrifuged at 35,000 rpm at 4°C for 5 h. Because the two peaks for the free and mRNA-bound 70S ribosomes are very close in mass and clean separation was not possible, the MATLAB file-exchange package, *peakfit*, was used to fit the two overlapped peaks as two Gaussian distributions. For the free ribosome control, cells were treated with 100μM puromycin for 5 minutes and collected. For serine hydroxamate (SHX, Sigma S4503) treatment, cells were grown in MOPS glucose minimal media until OD ~0.3 and treated with SHX with final concentration of 1 mg/mL for ten minutes before collection.

### *lacZ* induction and translational elongation rate measurement

The method was adapted from Zhu et al^47^. A final concentration of 5 mM IPTG (I2481C-25, Gold Biotechnology) was added to cultures. At every 15 seconds, 1 mL of culture was collected in a tube containing 10μL 100mM chloramphenicol, snap frozen in liquid nitrogen and stored at −20°C before subsequent measurement. After cells were thawed, 400μL of the sample was added to 100 μL 5x Z-buffer (0.3M Na_2_HPO_4_.7H_2_O, 0.2M NaH_2_PO_4_.H_2_O, 50mM KCl, 5mM MgSO_4_, 20mM β-mercaptoethanol) and incubated at 37°C for 10 minutes. 100μL of 4mg/mL MUG (337210010, ACROS Organics) in DMSO was then added to each sample every 10 seconds to accurately control the reaction time. The samples were incubated at 37°C in a thermomixer at 1,400 rpm mixing rate for 30 minutes to 2 hours, depending on the enzyme expression level. The reaction was stopped by addition of 300μL of 1M Na_2_CO_3_. The tubes were spun down at 16,000 X g for 3 minutes to sediment the cell debris. 200 μL of supernatant were taken and measured fluorescence by a microplate reader (365 nm excitation and 450 nm emission filter).

To infer translational elongation rate, the square root of the signal in excess of the signal at time zero was plotted. A linear fit was performed on the points after signal began to increase. The lag time is the x-intercept of the line. A previous study has shown that the initiation time remains constant for about 10 seconds across a wide range of conditions tested^47^. Therefore, we corrected the elongation times measured by subtracting 10 seconds in the lag times measured.

### Ribosome footprinting and total RNA extraction for RNA-Seq

The cell collection step was the same as for polysome profiling except that 1mM chloramphenicol was used in the sucrose solution. The footprinting and library preparation steps were adapted from Li *et al.*^2^ After quantification of RNA concentration with NanoDrop, samples with 500μg RNA were digested with 750U MNase (10107921001, Roche) for 1 hour at 25°C before being quenched with 6mM EGTA. The lysates were then layered onto a 10%-55% sucrose gradient and centrifuged. The monosome fraction was collected and snap frozen in liquid nitrogen. No polysome peaks were observed, indicating a thorough digestion. The RNA was isolated using hot phenol and size selected on 15% TBE-Urea PAGE gels run for 1 hour at 210V. Gels were stained with SYBR Gold and visualized using Dark Reader (Clare Chemical Research). RNA fragments with size between 25-40 nt were extracted using isopropanol precipitation. Total RNA was extracted with TRIZOL from the same pulverized cells used for footprinting. After DNaseI (04716728001,Roche) treatment and cleanup using RNA clean & concentrator 5 (R1016, Zymo Research), ribosomal RNA was subtracted using MicrobeExpress (AM1905, Ambion). The recovered RNA was fragmented and size selected. RNA fragments with size between 25-40nt were extracted using isopropanol precipitation.

### Library preparation and sequencing

Fragments from footprints and total RNA were dephosphorylated at the 3’ end by PNK (M0201, NEB). The repaired fragments were linked to the Universal miRNA Cloning Linker (S1315S, NEB), reverse transcribed (18080044, Thermo)and circularized (CL4111K, Epicentre). The circularized samples were PCR amplified (M0531L, NEB) and size selected. High quality PCR samples checked by Bioanalyzer high sensitive DNA chip. Deep sequencing was done by Illumina HiSeq 2500 on Rapid flowcells with settings of single end and 75 nt-long read length.

### Mapping and sequencing data analysis

Data manipulation including barcode splitting, linker trimming and mapping were done using Galaxy. The processed reads were mapped to *Escherichia coli* genome escherichia_coli_k12_nc_000913_3 from the NCBI database with the BWA short read mapping algorithm. Only the reads between 20-45 nt that aligned to the coding region were used for further analysis.

To infer the ribosome A-site position, python package Plastid^48^ was used to align the 3’ end of reads to the stop and start codons^49^, which are known to have higher ribosome densities. We found that the offsets were 12 nt for stop codon and 15 nt for start codon. Therefore, we used 11nt for A site position and 14nt for P site. The counts were normalized to the total counts in the coding region as reads per million (RPM) and transcripts per million (TPM). Further analysis was done using customized Python and R codes with packages including Plastid, dplyr, tidyr and ggplot2.

### Analysis of ribosome profiling data

Transcripts per million (TPM) was used to identify highly expressed genes for codon occupancy analysis. After assigning each mapped read to the A-site nucleotide, the raw counts were first normalized by the length of their mapped coding region, to yield count density for each gene. These count densities were then globally normalized to one million counts across all genes within one sample. To determine codon occupancy, transcripts having total counts over 100 TPM and containing more than 200 codons were considered. Ribosome footprint counts for the first and last 40 codons were removed to avoid possible effects from initiation and termination, and counts per codon were recorded for the remaining counts. For each gene and each codon type, the codon occupancy ratio is defined as the ratio of measured counts to expected counts. Expected counts are simply proportional to the frequency of that particular codon in the gene. The final reported codon occupancy ratio is the average of codon occupancy ratios from all genes considered weighted equally.

We calculated ribosome counts along a transcript using reads per million (RPM). Only transcripts having total counts over 10 TPM and containing more than 100 codons were considered. Total counts after filtering were normalized to one million reads. The selected genes account for more than 85% of the total counts. The ribosome counts are the sum of RPM at each position.

To do the first and second halves comparison of ribosome counts of genes, the same set of genes and trimming processing for codon occupancy were used. The RPM from positions were summed up based on its assigned location (first or second half) on the transcript and plot against each other.

## Code availability

All the codes used for data analysis in this paper are available upon request.

## Data availability

All the data used to reach the conclusion of this paper will be made available upon publication on SRA BioProject with submission number <u>SUB3256640</u>.

## References and Notes

1. You, C. et al. Coordination of bacterial proteome with metabolism by cyclic AMP signalling. 500, 301–306 (2013).

2. Li, G.-W., Burkhardt, D., Gross, C. & Weissman, J. S. Quantifying absolute protein synthesis rates reveals principles underlying allocation of cellular resources. Cell 157, 624–635 (2014).

3. Scott, M., Klumpp, S., Mateescu, E. M. & Hwa, T. Emergence of robust growth laws from optimal regulation of ribosome synthesis. Mol. Syst. Biol. 10, 747–747 (2014).

4. Giordano, N., Mairet, F., Gouzé, J.-L., Geiselmann, J. & de Jong, H. Dynamical Allocation of Cellular Resources as an Optimal Control Problem: Novel Insights into Microbial Growth Strategies. PLoS Comput. Biol. 12, e1004802 (2016).

5. Bremer, H. & Dennis, P. P. Modulation of Chemical Composition and Other Parameters of the Cell at Different Exponential Growth Rates. EcoSal 1–56 (2012).

6. Baracchini, E. & Bremer, H. Determination of synthesis rate and lifetime of bacterial mRNAs. Anal. Biochem. 167, 245–260 (1987).

7. Schaechter, M., Maaløe, O. & Kjeldgaard, N. O. Dependency on Medium and Temperature of Cell Size and Chemical Composition during Balanced Growth of Salmonella typhimurium. Microbiology 19, 592–606 (1958).

8. Scott, M., Gunderson, C. W., Mateescu, E. M., Zhang, Z. & Hwa, T. Interdependence of cell growth and gene expression: origins and consequences. Science 330, 1099–1102 (2010).

9. Yamamoto, T., Izumi, S. & Gekko, K. Mass spectrometry of hydrogen/deuterium exchange in 70S ribosomal proteins from E. coli. FEBS Lett. 580, 3638–3642 (2006).

10. Nomura, M. & Gourse, R. Regulation of the synthesis of ribosomes and ribosomal components. Annual review of … (1984).

11. Maaløe, O. & Kjeldgaard, N. O. Control of Macromolecular Synthesis, WA Beniamin. (Inc., 1966).

12. Dai, X. et al. Reduction of translating ribosomes enables Escherichia coli to maintain elongation rates during slow growth. Nat Microbiol 2, 16231 (2016).

13. Klumpp, S., Scott, M., Pedersen, S. & Hwa, T. Molecular crowding limits translation and cell growth. Proc. Natl. Acad. Sci. U.S.A. 110, 16754–16759 (2013).

14. Lewin, B. Genes VIII. (Pearson education, Inc, 2003).

15. Wanner, B. L. Gene regulation by phosphate in enteric bacteria. Journal of Cellular Biochemistry 51, 47–54 (1993).

16. Ahn, K. & Kornberg, A. Polyphosphate kinase from Escherichia coli. Purification and demonstration of a phosphoenzyme intermediate. J. Biol. Chem. 265, 11734–11739 (1990).

17. Schleif, R., Hess, W., Finkelstein, S. & Ellis, D. Induction kinetics of the L-arabinose operon of Escherichia coli. J. Bacteriol. 115, 9–14 (1973).

18. Dalbow, D. G. & Young, R. Synthesis time of β-galactosidase in Escherichia coli B/r as a function of growth rate. Biochemical Journal 150, 13–20 (1975).

19. Qin, D. & Fredrick, K. Analysis of polysomes from bacteria. Meth. Enzymol. 530, 159–172 (2013).

20. Beller, R. J. & Lubsen, N. H. Effect of polypeptide chain length on dissociation of ribosomal complexes. Biochemistry 11, 3271–3276 (1972).

21. Nathans, D. Puromycin inhibition of protein synthesis: incorporation of puromycin into peptide chains. Proc. Natl. Acad. Sci. U.S.A. 51, 585–592 (1964).

22. Steitz, J. A. Polypeptide Chain Initiation: Nucleotide Sequences of the Three Ribosomal Binding Sites in Bacteriophage R17 RNA. 224, 957–964 (1969).

23. Ingolia, N. T., Ghaemmaghami, S., Newman, J. R. S. & Weissman, J. S. Genome-wide analysis in vivo of translation with nucleotide resolution using ribosome profiling. Science 324, 218–223 (2009).

24. Doucette, C. D., Schwab, D. J., Wingreen, N. S. & Rabinowitz, J. D. α-Ketoglutarate coordinates carbon and nitrogen utilization via enzyme I inhibition. Nature Chemical Biology 7, 894–901 (2011).

25. Yuan, J. et al. Metabolomics-driven quantitative analysis of ammonia assimilation in E. coli. Mol. Syst. Biol. 5, 302 (2009).

26. Ikeda, T. P., Shauger, A. E. & Kustu, S. Salmonella typhimuriumApparently Perceives External Nitrogen Limitation as Internal Glutamine Limitation. Journal of Molecular Biology 259, 589–607 (1996).

27. Subramaniam, A. R., Pan, T. & Cluzel, P. Environmental perturbations lift the degeneracy of the genetic code to regulate protein levels in bacteria. Proc. Natl. Acad. Sci. U.S.A. 110, 2419–2424 (2013).

28. Sørensen, M. A. et al. Over Expression of a tRNALeu Isoacceptor Changes Charging Pattern of Leucine tRNAs and Reveals New Codon Reading. Journal of Molecular Biology 354, 16–24 (2005).

29. Elf, J., Nilsson, D., Tenson, T. & Ehrenberg, M. Selective charging of tRNA isoacceptors explains patterns of codon usage. Science 300, 1718–1722 (2003).

30. Hauryliuk, V., Atkinson, G. C., Murakami, K. S., Tenson, T. & Gerdes, K. Recent functional insights into the role of (p)ppGpp in bacterial physiology. Nat Rev Micro 13, 298–309 (2015).

31. Potrykus, K. & Cashel, M. (p)ppGpp: Still Magical?*. Annu. Rev. Microbiol. 62, 35–51 (2008).

32. Mitkevich, V. A. et al. Thermodynamic characterization of ppGpp binding to EF-G or IF2 and of initiator tRNA binding to free IF2 in the presence of GDP, GTP, or ppGpp. Journal of Molecular Biology 402, 838–846 (2010).

33. Milon, P. et al. The nucleotide-binding site of bacterial translation initiation factor 2 (IF2) as a metabolic sensor. Proc. Natl. Acad. Sci. U.S.A. 103, 13962–13967 (2006).

34. Jiang, M., Sullivan, S. M., Wout, P. K. & Maddock, J. R. G-protein control of the ribosome-associated stress response protein SpoT. J. Bacteriol. 189, 6140–6147 (2007).

35. Gentry, D. R. & Cashel, M. Mutational analysis of the Escherichia coli spoT gene identifies distinct but overlapping regions involved in ppGpp synthesis and degradation. Mol. Microbiol. 19, 1373–1384 (1996).

36. Pavlov, M. Y. & Ehrenberg, M. Optimal control of gene expression for fast proteome adaptation to environmental change. Proc. Natl. Acad. Sci. U.S.A. 110, 20527–20532 (2013).

37. Kjeldgaard, N. O., Maaløe, O. & Schaechter, M. The Transition Between Different Physiological States During Balanced Growth of Salmonella typhimurium. Microbiology 19, 607–616 (1958).

38. Brauer, M. J. et al. Coordination of growth rate, cell cycle, stress response, and metabolic activity in yeast. Mol. Biol. Cell 19, 352–367 (2008).

39. Paul, B. J., Ross, W., Gaal, T. & Gourse, R. L. rRNA Transcription in Escherichia coli. Annu. Rev. Genet. 38, 749–770 (2004).

40. Laursen, B. S., Sørensen, H. P., Mortensen, K. K. & Sperling-Petersen, H. U. Initiation of protein synthesis in bacteria. Microbiology and Molecular Biology Reviews 69, 101–123 (2005).

41. Rojas, A.-M., Ehrenberg, M. N., Andersson, S. G. E. & Kurland, C. G. ppGpp inhibition of elongation factors Tu, G and Ts during polypeptide synthesis. Mol. Gen. Genet. 197, 36–45 (1984).

42. Zarrinpar, A., Chaix, A., Yooseph, S. & Panda, S. Diet and Feeding Pattern Affect the Diurnal Dynamics of the Gut Microbiome. Cell Metabolism 20, 1006–1017 (2014).

43. Fischer, E. & Sauer, U. Large-scale in vivo flux analysis shows rigidity and suboptimal performance of Bacillus subtilis metabolism. Nat. Genet. 37, 636–640 (2005).

44. Towbin, B. D. et al. Optimality and sub-optimality in a bacterial growth law. Nat Commun 8, 14123 (2017).

45. Peters, J. M. et al. A Comprehensive, CRISPR-based Functional Analysis of Essential Genes in Bacteria. Cell 165, 1493–1506 (2016).

46. Baba, T. et al. Construction of Escherichia coli K-12 in-frame, single-gene knockout mutants: the Keio collection. Mol. Syst. Biol. 2, 2006.0008 (2006).

47. Zhu, M., Dai, X. & Wang, Y.-P. Real time determination of bacterial in vivoribosome translation elongation speed based on LacZα complementation system. Nucleic Acids Res. gkw698 (2016). doi:10.1093/nar/gkw698

48. Dunn, J. G. & Weissman, J. S. Plastid: nucleotide-resolution analysis of next-generation sequencing and genomics data. BMC Genomics 17, 958 (2016).

49. Woolstenhulme, C. J., Guydosh, N. R., Green, R. & Buskirk, A. R. High-precision analysis of translational pausing by ribosome profiling in bacteria lacking EFP. Cell Rep 11, 13–21 (2015).

